# Connectome spectrum electromagnetic tomography: a method to reconstruct electrical brain source-networks at high-spatial resolution

**DOI:** 10.1101/2022.07.26.501544

**Authors:** Joan Rué-Queralt, Hugo Fluhr, Sebastien Tourbier, Yasser Aleman-Gómez, David Pascucci, Jérôme Yerly, Katharina Glomb, Gijs Plomp, Patric Hagmann

## Abstract

Connectome Spectrum Electromagnetic Tomography (CSET) combines diffusion MRI-derived structural connectivity data with well-established graph signal processing tools to solve the M/EEG inverse problem. Using simulated EEG signals from fMRI responses, and two EEG datasets on visual-evoked potentials, we provide evidence supporting that (i) CSET captures realistic neurophysiological patterns with better accuracy than state-of-the-art methods, (ii) CSET can reconstruct brain responses more accurately and with more robustness to intrinsic noise in the EEG signal. These results demonstrate that CSET offers high spatio-temporal accuracy, enabling neuroscientists to extend their research beyond the current limitations of low sampling frequency in functional MRI and the poor spatial resolution of M/EEG.

## 1 INTRODUCTION

The human brain is a network of deeply interconnected neurons and complex architecture. Understanding its dynamic functioning at a sub-second level is of critical importance for several fields in medicine and technology [1, 2, 3, 4]. Neuroimaging techniques have advanced to a point where functional mapping of whole-brain activity in small animal models is now possible at sub-second level, thanks to techniques like calcium imaging [5, 6, 7]. However, human neuroimaging poses other challenges, as no modality offers both high spatial and temporal resolution [8]. Modalities with high spatial resolution, such as functional Magnetic Resonance Imaging (fMRI) or Positron Emission Tomography (PET), have lower temporal resolution compared to techniques such as magneto/electro-encephalography (M/EEG), which have higher temporal resolution but suffer from low or very poor spatial accuracy.

To address this limitation, Electromagnetic Tomography (ET), also known as electrical or M/EEG source imaging (ESI), has been proposed as a computational approach to estimate electrical neuronal activity at the whole-brain scale. By combining M/EEG recordings and structural MR images, ET aims to achieve better spatial resolution while maintaining fast sampling of neural signals [9] (Fig. 1a). ET has allowed significant advances in several fields of brain functional mapping, including epilepsy [10], sleep [11], cognition [12], and brain-computer interfaces [13].

**FIGURE 1.**
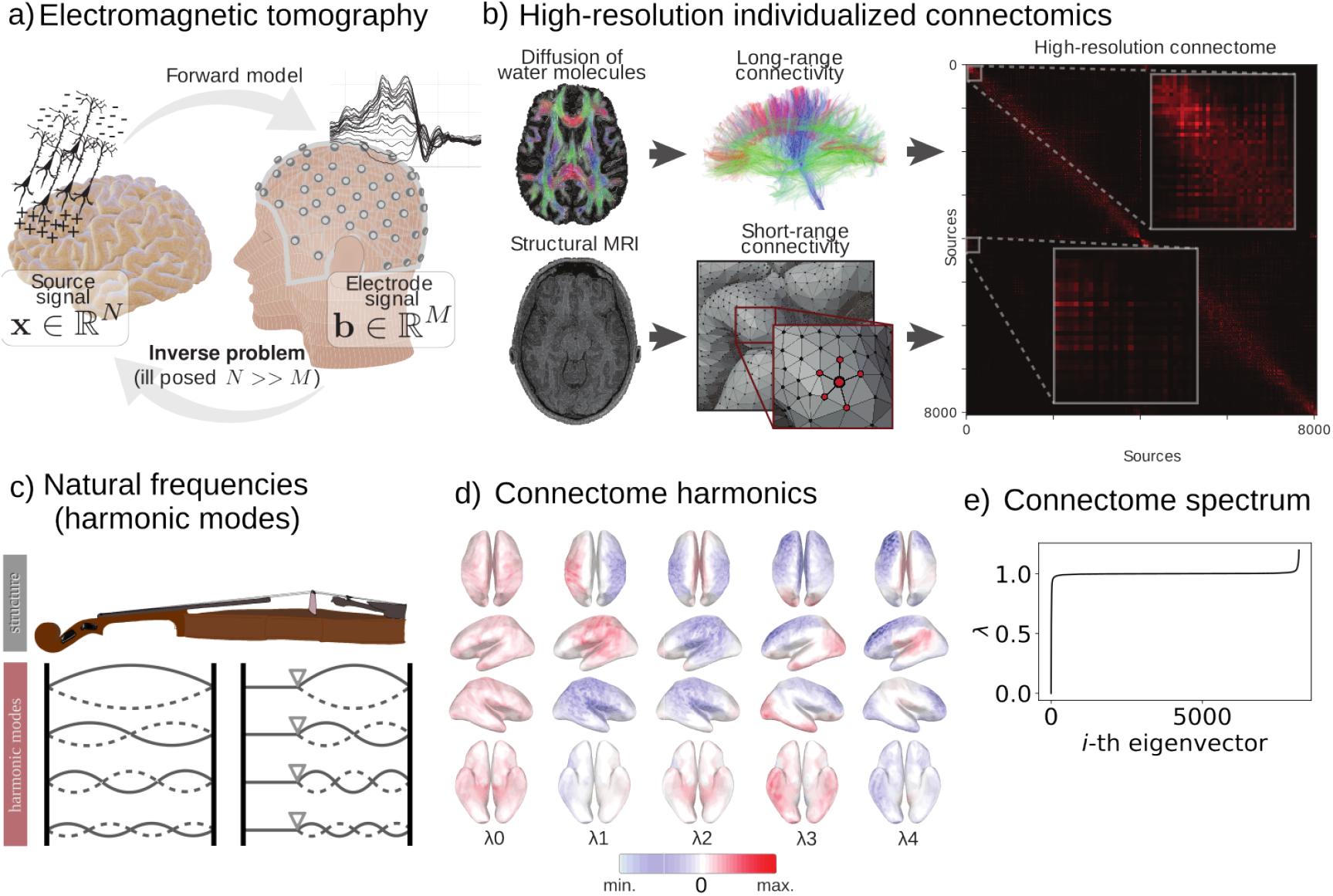
Connectome Spectrum Electromagnetic Tomography (CSET) pipeline. a) Illustration of the EEG inverse problem. The EEG inverse problem is the process of estimating of the electrical sources in the brain that generated the measured electrode signal at the scalp. This problem is ill-posed because the number of sources (N) is much larger than the number of electrodes (M). There exist an infinite number of source combinations that could generate a single electrode signal. For this reason, mathematical regularization based on mechanistic or empirical assumptions of brain activity are needed. b) The high-resolution individualized connectomes are constructed by combining the long-range white-matter connectivity (based on MR tractography) and the short-range cortical connectivity (based on Euclidean distance). c) The harmonic modes of a physical object refer to the different ways an object can resonate when a force is applied. In the case of a violin, the harmonic modes or natural frequencies, are typically determined by the length of the string and its tension. d) The harmonic modes of the high-resolution connectome are determined by the connectivity pattern, they also have a notion of frequency, or smoothness. Connectome harmonics are identified as the eigenvectors of the high-resolution connectome graph Laplacian. Here only the smoothest five eigenvectors are shown. e) The connectome spectrum, i.e., the eigenvalues corresponding to the connectome harmonics, which indicate how smooth each harmonic is.

Electromagnetic tomography focuses on solving two different processes or problems: the forward problem and the inverse problem. The **forward problem** involves determining the relationship between the effective electric sources in the brain and the measurements recorded by scalp electrodes or magnetometers. Specifically, it describes the propagation of electric fields in the head (as seen in Fig. 1a), from the electrical depolarization of pyramidal cells in the cortex to the M/EEG sensors. This problem is solved using Maxwell’s equations, which take into account the different conductivity properties of the brain tissue [14].

Solving the forward problem accurately requires discretizing the head volume and mapping realistic tissue conductivity values in it, which can be challenging. However, there are solid implementations available in the literature that rely on finite element modeling and personalized mappings based on individual brain MRI scans [14]. These solutions have been shown to be effective in addressing this challenge.

The **inverse problem**, on the other hand, is the mathematical formulation of the tomography itself. It involves modeling the neuronal activity as a function of the measurements recorded by scalp electrodes or magnetometers. By combining the forward and inverse problems, ET attempts to estimate the electrical neuronal activity across the entire brain with high spatial and temporal resolution.

Each sensor in M/EEG recordings captures activity from multiple sources within the brain’s gray matter, and the source electric fields propagate through the tissues non-uniformly, depending on local conductivities and morphology. The inverse problem of discerning which sources, or combination of sources, are responsible for a given M/EEG measurement is ill-posed, meaning there is no unique solution for the problem [14]. There are infinitely many solutions that can be consistent with the measured M/EEG data. Thus, regularization is needed to solve the inverse problem, which involves making strong assumptions about the spatial distribution of the sources [9].

However, the current models used to solve the M/EEG inverse problem are known to produce low-resolution reconstructions [15, 16, 17, 18, 19] or make unrealistic assumptions [20]. These limitations pose significant challenges for accurately mapping electrical brain activity, and there is a pressing need for biologically plausible and realistic models to overcome these hurdles. Such models would be able to encompass a wide range of neural activation patterns, making it possible to accurately solve the inverse problem and improve our understanding of brain function.

Over the past two decades, a growing body of evidence has demonstrated that the patterns of human brain activity are tightly constrained by the underlying structural connectivity [21, 22, 23, 24, 25, 26]. It is widely recognized that taking into consideration brain structural connectivity when analysing brain activity signal is of crucial importance for proper interpretation [27, 28, 29, 30]. It seems that similar to any other physical object, such as a metal plate or a vibrating rope, the resonant frequencies of the brain are largely determined by its underlying structure [31]. Recent data suggest that brain activity can be efficiently represented as a combination of its normal modes [32], which are known as connectome harmonics and form the building blocks of well-known brain functional networks associated with both rest and different tasks (Fig. 1c,d). The representation of brain activity in the basis defined by connectome harmonics, also known as connectome spectral representation, is the result of a graph Fourier transform [33] on the connectome, which involves the eigen-decomposition of the graph Laplacian. Studies have found that brain activity during visual perception [34, 35] is characterized by a sparse connectome harmonic representation, indicating that the connectome harmonics provide a powerful framework for understanding the functional organization of the brain. Similarly, other studies have found similar properties while investigating different states of consciousness [36].

In this work, we introduce a new approach, called Connectome Spectrum Electromagnetic Tomography (CSET), which aims to solve the M/EEG inverse problem by taking advantage of the sparsity of brain activity in its connectome spectral representation. Specifically, we model this property as a prior probability of the sources (Fig. 1c-f), within a Bayesian optimization framework (very similar in essence to the compressed sensing framework [37, 38]. While prior methods have used sparsity priors at the source domain [39, 40, 41], these are limited to highly localized neural activity and do not capture distributed neural networks. The underlying assumptions of CSET make it better suited for reconstructing distributed brain activity.

To evaluate the effectiveness of CSET, we applied it to both simulated EEG data from brain activity patterns corresponding to fMRI task activation (Fig. 2) and real EEG data from two experiments on visual evoked potentials (Fig. 3). Our results demonstrate that incorporating the high-resolution connectivity structure of the brain (Fig. 1b) improves the signal-to-noise ratio (SNR) and the accuracy of the reconstructed brain activity compared to state-of-the-art approaches that do not use this information.

**FIGURE 2.**
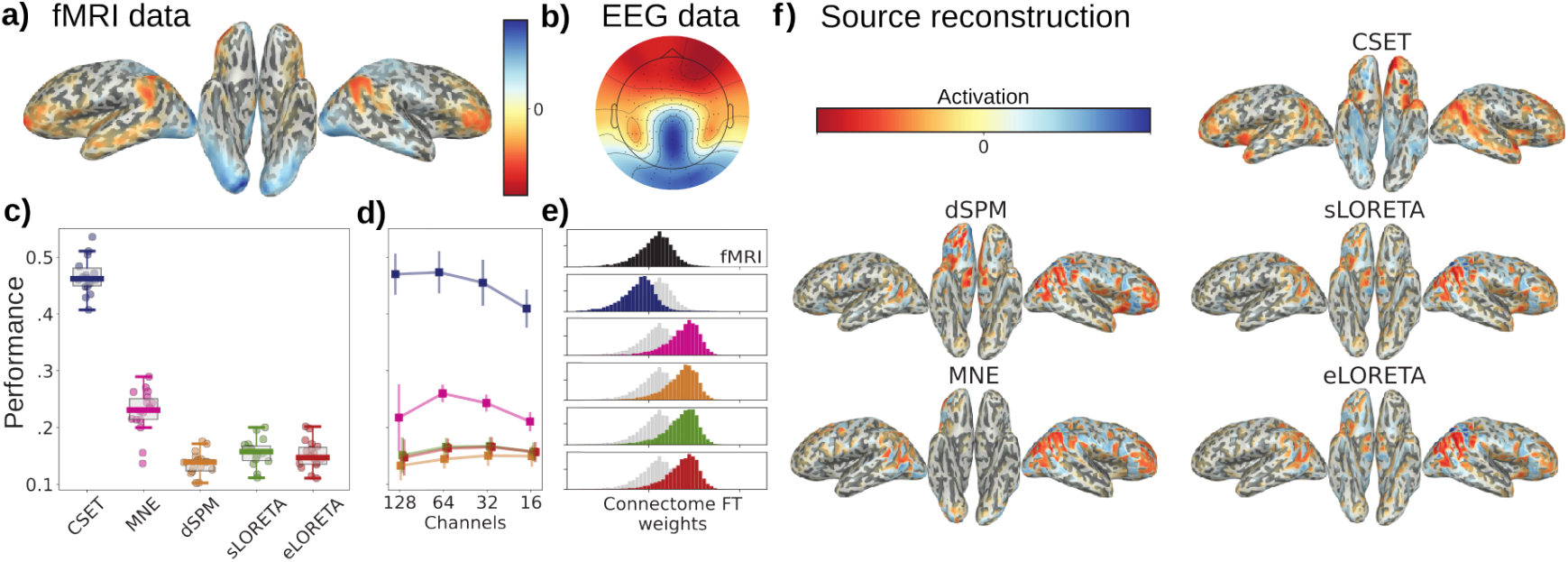
Results of simulation experiment. a) The ground truth signal used for simulating the EEG data corresponded to the fMRI response of each subject to a face perception task. b) The simulated EEG is obtained by applying the forward model to the ground truth signal, i.e., propagating the source activation to the electrodes through the lead-field matrix. c) The performances (*r*^2^ between ground truth and reconstructed signal) of the different reconstructions when using the full montage of 128 electrodes. d) The performances of the different reconstructions when using the full and sub-sampled montages. e) Distributions of the connectome Fourier transform weights, i.e., the resulting coefficients of applying the Fourier transform on the source reconstructed signal, for the ground truth data and the different reconstructions. f) Source reconstructions for each method for the same subject.

**FIGURE 3.**
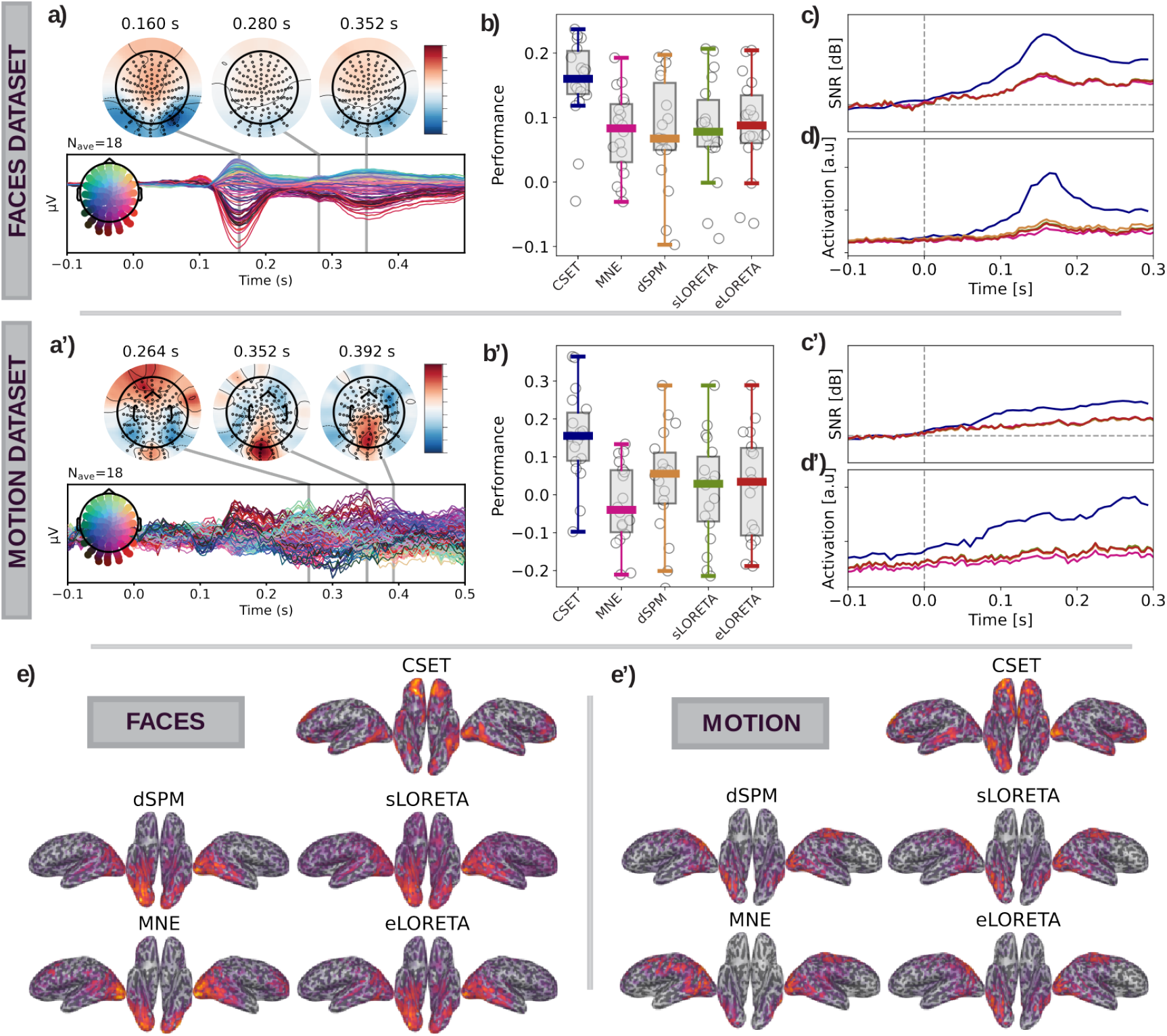
Results of visual evoked potentials. a-a’) Residual visual evoked potentials (a): The EEG response to faces minus the response to scrambled faces; (a’): The EEG response to coherent motion minus the response to random motion. The temporal traces correspond to the difference in EEG measured potential evoked by the experimental condition and the control. The topographic maps corresponding to the measured data at the peak-times of the traces are plotted on top. b, b’) The performances of the different reconstruction methods for ventral (b), and dorsal (b’) visual systems (performance: *r*^2^ between reconstructed signal and fMRI signal from an equivalent task). c-c’) Signal-to-noise ratio (SNR) dynamics for different reconstructions for pre- and post-stimulus presentation time (SNR defined as power of post stimulus vs power pre-stimulus). d-d’) Region of interest dynamics: right fusiform gyrus (FFA) for the face perception task; and the bilateral posterior middle-temporal area (MT area) for the motion perception task. e-e’) Group average (mean) reconstructions for the different source reconstruction methods.

## 2 MATERIALS AND METHODS

### 2.1 The high-resolution structural connectome

The structural connectome defines how the axonal fiber bundles in the brain’s white matter support the connectivity between different gray matter regions. For the VEPCON dataset, we estimated the tractograms from the diffusion weighted imaging (DWI) data of each participant, using Connectome Mapper 3 [42]. The tractography algorithm was performed after denoising the diffusion data with MRTRIX MP-PCA, bias field correction with ANTS-N4, eddy current and motion correction from FSL. For each subject we launched 10 million deterministic streamlines (with ACT) in the white matter, which were posteriorly filtered with SIFT. After that, we followed the approach presented by Mansour and colleagues [43] to obtain high-resolution individual connectomes at the resolution of approximately 8000 nodes on the brain’s gray matter surface. To allow other scientists to use our source imaging framework on data sets with no available diffusion MRI data, we constructed a group consensus connectome from the high-resolution individual connectomes of the *VEPCON dataset*. The consensus connectome was created based on the distance-dependent thresholding method [44].

### 2.2 Connectome harmonics

The structural connectome defines a graph object 𝒢 (𝒩, ε), in which the nodes 𝒩 of the graph represent different regions in the brain cortical surface, and the edges E of this graph describe the connectivity strength between each pair of nodes as estimated from the high-resolution connectome. Analogous to conventional signal processing spectral analysis, graph signal processing allows us to study the signal in terms of its graph (i.e. spatial) frequency content (for an in depth review of graph-signal processing, see [33]).

To obtain the connectome spectrum of the brain activity signal we first need to perform an eigendecomposition of the normalized connectome graph Laplacian:

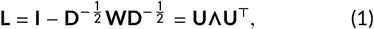

where **I** is the identity matrix, **W** is the connectome graph’s weight matrix (here defined as the number of streamlines connecting each pair of brain reagions) and **D** is the degree matrix (i.e., a diagonal matrix with the degree value of each node as the diagonal elements). **U** is a matrix that contains the eigenvectors of the graph Laplacian in its columns, defining the connectome graph Fourier basis, and **Λ** is a diagonal matrix contains its eigenvalues *λ*_*i*_, which are associated to the notion of frequency in traditional signal processing. The connectome spectrum 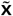 of the brain activity signal **x** is thus obtained by means of the graph Fourier transform

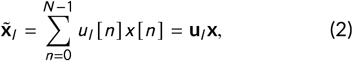

where *l* indexes the connectome harmonics (i.e. eigenvectors) and *n* the nodes. The reader is referred to our previous publications for further details on this representation of the brain activity signal [34, 35].

### 2.3 The forward model

The forward model of the M/EEG imaging system describes how the magnetic/electrical currents are propagated from their origin in the active neurons residing in the gray-matter towards the recording sensors in the scalp. When a pyramidal neuron in the gray-matter receives an excitatory postsynaptic potential (EPSP), its voltage dependent sodium channels open, the positively charged sodium ions enter in the neuron and due to the electrical neutrality conservation principle, an active source of current is produced in the apical region of that neuron. This creates an electrical dipole. When many neighbouring pyramidal neurons activate simultaneously, they generate an electrical dipole that is strong enough to traverse the different tissues of the head, and to be measured by M/EEG electrodes [14]. The forward model characterizes the probability of the M/EEG measurements on the scalp *p* (**b**_*t*_ |**x**_*t*_) for **b**_*t*_ ∈ ℝ^*M*^, conditioned **x**_*t*_ ∈ ℝ^*N*^ being the true cortical source activity (*M* being the number of measuring sensors and *N* the number of neuronal activity sources). The M/EEG system deals with noise that is due to independent random perturbations at the sensor level, following a zero-mean Gaussian distribution [14]:

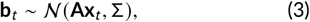

where **A** ∈ ℝ^*M* ×*N*^ is known as the lead-field matrix, and its components *A*_*i,j*_ define the contribution of the *j* -th cortical source to the *i* -th M/EEG scalp sensor, which are estimated by solving Maxwell’s equations [14]. To model independent and identically distributed noise across sensors we use Σ = **I***σ*. Given our primary focus on the inverse problem, we rely on the default forward computation pipeline implemented in *MNE-Python* [45], as it is a well-documented method in an openly available software toolbox. In particular, we used the default MNE-Python surface-based boundary element method (BEM) approach [46], where the boundary surfaces are tessellated into a mesh of triangles with different conductivity values [inner-skull = 0.3, outer-skull = 0.006, outer-skin-skull = 0.3].

In this work, we constrain the sources to be normally oriented to the cortical surface of the brain. There exist two main reasons behind this choice. First, the complexity of the algorithm is reduced by reconstructing a single value per source rather than a value per each coordinate axis (*x, y* and *z*). Second, the dipoles originated due to the excitation of pyramidal neurons are mostly oriented normally to the cortical surface [14].

### 2.4 The inverse problem

From a Bayesian perspective, the inverse problem can formulated as trying to find the most likely electrical source configuration 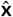 in the gray-matter given the scalp measurements **b**, the forward model **A** and a prior probability over the distribution of x. This is known as the maximum a posterior (MAP) estimate:

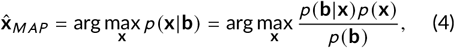

where *p* (**x|b**) is the posterior probability, *p* (**b|x**) is the likelihood function, and *p* (**x**) defines the prior. *p* (**b**) does not affect the maximizer argument and can be ignored. In practice, we simplify the optimization problem by taking the negative log-transform of the posterior distribution, so that we minimize over a sum instead of maximize over a product:

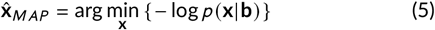

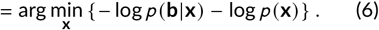

Given our assumption of normally distributed EEG data (Eq. 3), the likelihood function of the data can be defined by a Gaussian distribution:

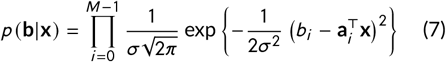

and its negative log-transform results in:

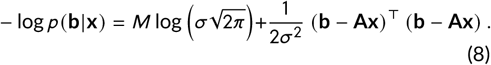

This results in a likelihood function with a quadratic term plus some constant term. The constant term does not affect the argument of the minimization problem and can be ignored, thus leaving with the well-known least-squares form:

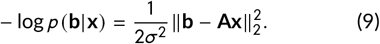

For the prior distribution, we assume that brain activity follows a sparse connectome spectral representation, which is well modeled by the Laplacian probability distribution on the graph-Fourier transformed coefficients:

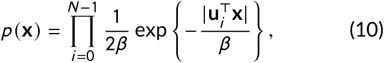

where *β* is a scale parameter related to the variance of the distribution. Its negative log-transform results in:

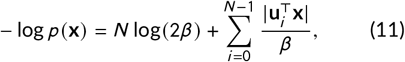

which, ignoring the constant term, results in the *L*1-norm:

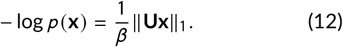

Combining Eqs. 9 and 12, we can rewrite Eq. 5 as:

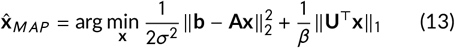

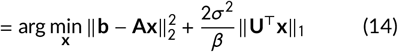

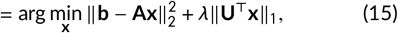

where we have used the regularization parameter 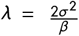 defines the uncertainty trade-off between the likelihood and the prior. The *L*1-norm acts as a regularization function that imposes structure (sparsity on some transform F). Another common way to look at this problem is the following. Given that the number of gray-matter sources *N* is much larger than the number of M/EEG sensors *M*, the inverse problem is under-determined or ill-posed. This means that there exist an infinite number of source activity configurations (x) that can produce the measured EEG scalp potential (b) at a given time-point. The inclusion of a regularization function constraints the number of solutions to a single one.

### 2.5 Connectome Spectrum Electrical Tomography (CSET)

We have shown in previous publications that the neuronal activity is sparsely represented by the connectome-based graph Fourier transform (see *Connectome Spectral Analysis*), which decomposes brain activity into a small number of active brain networks [35, 34]. In addition to our research, other studies have established a theoretical basis for the structural constraint on functional connectivity [32, 36, 47, 48, 49]. These works have revealed that neural mechanisms relying on delayed excitatory-inhibitory interactions facilitate the self-organization towards exciting a relevant set of eigenmodes [32, 48]. Moreover, previous investigations have successfully applied structural eigenmodes to explain brain activity during rest at very short timescale [49] and evoked activity [47]. These findings suggest that these eigenmodes play a crucial role in dynamically integrating and segregating information across the cortex, thus serving important cognitive functions.

In this study, we present Connectome Spectrum Electromagnetic Tomography (CSET), a novel approach that addresses the optimization problem outlined in Eq. 13. We achieve this by representing brain activity as a constrained combination of active eigenmodes, emphasizing sparsity within the connectome spectrum basis:

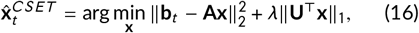

where 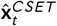 is the CSET source reconstruction at time *t*, **b**_*t*_ is the EEG data vector at time *t*, **U** is the connectome-based graph Fourier transform matrix containing the approximately 8000 eigenvectors of the structural connectivity normalized graph Laplacian. The following steps are performed by CSET to reconstruct sources:

#### 1. Depth normalization

The least squares term in Eq. 16 term is known to bias the optimized solution towards sources that are closer to the electrodes [50]. This can be alleviated by incorporating adding penalty function with a weighting factor composed by an *L*_2_-norm term [51]. Here, in order to avoid the increased computational effort of this approach, we adopt a strategy that is also implemented in *MNE-Python*, namely weighting the rows of the leadfield matrix prior to reconstruction:

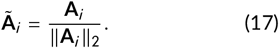

#### 2. M/EEG data normalization

The statistics of the measured signal will affect the optimal parameter *λ* in Eq. 16. To make the choice of *λ* not rely on the measurements, we normalize the measured signal as follows:

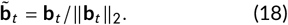

#### 3. Minimization via accelerated proximal gradient descent of

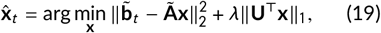

Eq. 19 contains a non-differentiable functional (*L*_1_-norm) and iterative optimization methods based solely on the gradient, such as gradient descent, cannot be used. Instead, we relied on the accelerated proximal gradient descent method (APGD) [52], a well-known primal-dual splitting optimization algorithm. The steps to reconstruct the sources from a single EEG time point using this algorithm are explained in Algorithm 1.

##### Algorithm 1 CSET

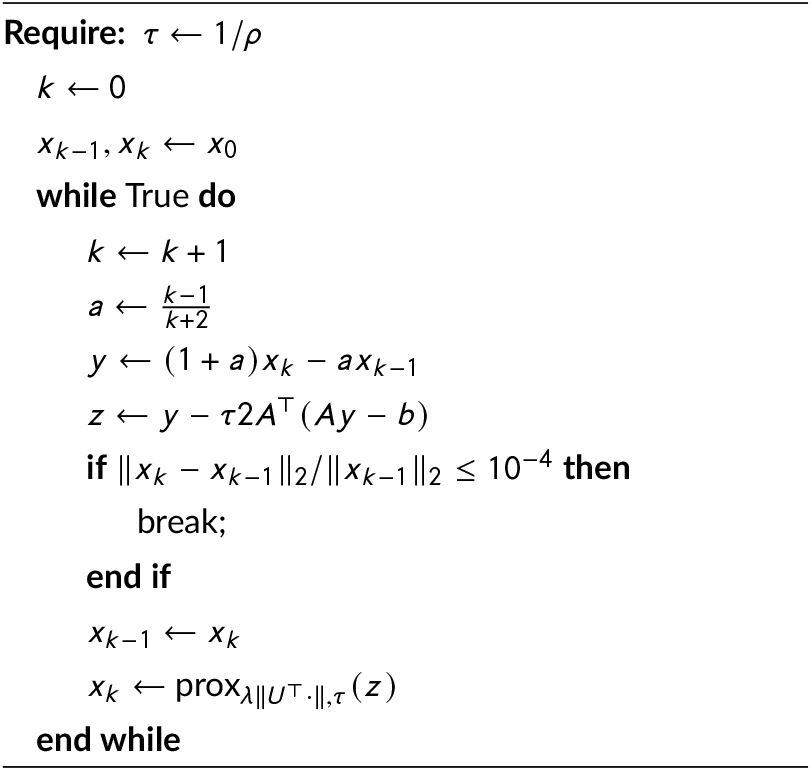

The optimization is solved with Pycsou [53], a Python package for solving linear inverse problems. The step *τ* = 1/*ρ* is selected with *ρ* being the gradient Lipschitz constant of the least squares term in Eq. 19, to fulfill the convergence rates guarantees. As a stopping criterion we selected an absolute relative error of *ϵ* = 10^−4^. The proximity operator is a mathematical tool that executes a step analogous to gradient descent but is specifically designed for non-smooth functions, such as those involving the *L*_1_-norm:

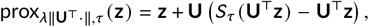

where, given an input vector z and a threshold *τ*, the soft thresholding operator is defined as:

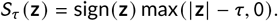

The soft thresholding operator has the effect of setting values of **x** with magnitude less than or equal to *λ* to zero, and otherwise shifting values of **x** toward zero by *λ*.

#### 4. Re-scale reconstruction

As a final step, the normalization factors obtained in steps 1 and 2 are rescaled back to the solution to the optimization problem, resulting in the final electrical source recon-struction:

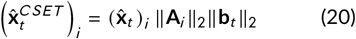

### 2.6 Datasets

Recently, two multimodal neuroimaging datasets have been released that have great potential to help develop source imaging tools as well as validate their performance and clinical relevance (*VEPCON* datasets from now on) [54]. In these datasets, visual evoked potentials were recorded for 20 participants while they discriminated between either briefly presented faces and scrambled faces on one hand, or coherently moving stimuli and incoherent ones on the other hand. This dataset is openly accessible (https://openneuro.org/git/0/ds003505). Apart from high-density EEG of visual evoked potentials, the datasets also include structural MRI, diffusion weighted images (DWI) and singletrial behavior. This allowed us to study the reconstruction of brain activity maps that activate in response to very well studied paradigms.

Apart from the data released along the original publication, here we also include fMRI spatial maps of the same participants, under a similar experimental task. These spatial maps are used in this work for two different purposes. In a first instance, for each subject of the VEPCON datasets, we used the fMRI activation pattern corresponding to the subjects’ response to a face stimulation paradigm as ground truth. This ground truth signal was used as simulated electrical brain activity, and then used to compare against the reconstruction from the simulated measurements.

Although neurovascular coupling—the complex mechanism that connects neural activity to the blood flow changes captured by fMRI—is not fully elucidated [55, 56], a significant body of evidence supports the notion that both fMRI and EEG signals are predominantly indicative of synaptic activity within the gray matter [57, 58]. Several studies have revealed a meaningful correlation between the fMRI signals (specifically, the BOLD response) and local field potentials (LFP), across an extensive frequency spectrum [59]. These findings intimate a shared origin of cortical synaptic activity between EEG and fMRI [60, 61], thereby highlighting their intertwined nature. Consequently, although EEG and fMRI signals manifest differences in spatial and temporal dimensions and are characterized by individual sensitivity, resolution, and specificity scales [62], a meticulous approach allows for a comparative analysis between the sources of fMRI and the estimates of EEG source reconstruction [63].

#### 2.6.1 MRI pre-processing

The acquisition of functional MRI data (fMRI) was performed at the HFR Fribourg – Hôpital cantonal (Fribourg, CH), using a Discovery MR750 3.0T (GE Healthcare, Chicago, USA). fMRI data were acquired while each subject performed two different visual tasks, one on faces, and another on moving dots.

Across subjects, the order of the two tasks was counterbalanced. For each task, we used a block structure similar to the one adopted for the EEG session (see Methods), but with 48 trials for each task condition (instead of 200). At the end of each block, the instruction to “REST” was presented, followed by a fixed break of 12s (Rest period). Exceptionally, the last Rest period that had a duration of 60s. In each trial, subjects had a limited time of 1,500ms to respond, after which their response was considered incorrect.

An MRI-compatible fiber optic response pad (Current Designs Inc., Philadelphia, USA) was used to collect the participant responses. The visual stimuli were presented on a NordicNeuroLab (Bergen, NO) MRI-compatible LCD monitor (32 inches diagonal size, 1920x1080 resolution, 405 c/m2 surface luminance, 4000:1 contrast, 60 Hz refresh rate, 6.5 ms response time), placed above the scanner-bed at 244 cm from the subject’s eyes and made visible to the subject through a mirror placed on the head coil. Those subjects who suffered from some eye disorder (e.g., myopia) wore MRI-compatible glasses with appropriate lenses for optical correction. The fMRI data were acquired using a T2*-weighted EPI sequence with 40 slices each, with slice-thickness of 3 mm, between-slices spacing of 0.3 mm, interleaved bottom-up slice acquisition, anterior-to-posterior phase encoding direction, repetition time (TR) of 2,500 ms, echo time (TE) of 30 ms, and flip-angle of 85°. The first 4 volumes of each run were discarded.

The Statistical Parametric Mapping (SPM) toolbox was used for preprocessing the fMRI data (toolbox version SPM12; University College London, UK; https://www.fil.ion.ucl.ac.uk/spm/). First, the functional images were aligned to the mean of each session, using a two-pass realignment procedure for motion correction, and then a slice-timing correction was applied 35. After realignment, the mean functional image was co-registered to the anatomical image using the normalized mutual information as the cost function. The SPM12 standard segmentation procedure was adopted to obtain the masks for cerebrospinal fluid (CSF) and white matter (WM), which were used to extract the time courses of CSF and WM signals for each subject. All images were then normalized to the Montreal Neurological Institute and Hospital (MNI) stereotaxic space with a fourth-order B-spline interpolation, and smoothed with a Gaussian filter with 8 mm FWHM kernel.

#### 2.6.2 fMRI statistical maps

A two-stage approach based on a general linear model (GLM) was employed to analyze the functional images. In this approach, the first-level analysis was implemented using a block design with two regressors of interest, each modeled with a boxcar function convolved with the canonical hemodynamic response function (HRF). Two regressors of interest were defined to model the two stimuli conditions (Faces vs. Scrambled, on vs. off motion of the disk). The GLM included also a set of nuisance regressors that modeled the six motion realignment parameters, the mean signals in CSF and WM, and a constant term. Finally, a high-pass filter (200 s cutoff) was applied to the functional images time series, which allowed removing noise at very low frequencies. The second-level group analysis was implemented on the previously obtained statistical maps and involved voxel-wise t-test comparison across participants. The obtained volumetric statistical maps were mapped to the same surface used for source reconstruction in the native space, using the *Freesurfer’s mri_vol2surf* function. A group estimate of the response pattern to the stimuli was estimated by morphing each surface map to the same space (subject “sub-01” native space). The average map was finally morphed back to the individual’
ss native space and used to assess the performance of source reconstruction.

### 2.7 Simulation of EEG data from fMRI Data

To simulate the EEG signal, we used the entire fMRI map, incorporating both positive and negative BOLD responses. The fMRI pattern was first standardized by dividing each voxel value by the standard deviation of the entire fMRI map. Subsequently, the pattern was shifted by its minimum value to exclude any negative values. We then applied the forward model with a matrix-vector multiplication with the lead-field matrix. To mimic realistic conditions, the generated EEG signals were corrupted with Gaussian noise. The noise power was calculated based on a predefined Signal-to-Noise Ratio (SNR) of 3 decibels (dB) as follows. While the spatio-temporal evolution of the neural sources is an important consideration, it was not explicitly modeled in this study for the sake of simplification and we limited our analysis to a single time point. Future studies are planned to incorporate the temporal dynamics in the simulation.

## 3 RESULTS

### 3.1 Increased sensitivity towards physiological patterns

*In-silico* simulations are a versatile tool for testing source reconstruction algorithms under various conditions, including different parameter configurations, data sampling strategies, and assumptions underlying the generative model of the signal. To minimize bias associated with selecting the generative model and to generate brain activity signals that are physiologically plausible, we used fMRI data as ground truth signals (Fig. 2a). Specifically, we utilized task fMRI to obtain a detailed spatial response to faces and moving dots for each participant. Fig. 2a shows the activation pattern in response to faces: the Fusiform Gyrus (FFA) is activated with some slight right dominance while the default-mode network is suppressed, as it is expected during the performance of a global visual discrimination task [64].

We evaluated the performance of our proposed method, Connectome Spectrum Electromagnetic Tomography (CSET), and four state-of-the-art methods available in *MNE-Python*, Minimum Norm Estimate (MNE), dynamic Statistical Parametric Mapping (dSPM [18]), and exact and standardized LOw Resolution Electric TomogrAphy (sLORETA [17], eLORETA [19]), by computing the r-squared metric, i.e., the square of the Pearson correlation coefficient, between the ground truth and reconstructed signals. These tools use the EEG signal covariance in their optimization algorithm, which we computed for pre-stimulus time (−200ms, 0ms) with the *MNE-Python* function *compute_covariance* [65], with the methods “shrunk” and “empirical”. In the determination of the regularization parameter *λ*, we employed a logarithmic scale spanning 20 distinct values, ranging from 10^−5^ to 0.3. Our analysis showed that CSET, which utilizes the subject specific brain connectivity map to reconstruct brain activity, outperformed the other methods in terms of reconstruction accuracy under different electrode montages (Fig. 2c-d). In fact, CSET recovered the ground truth signal with over twice the accuracy of the second-best method (r = 0.46 versus r = 0.23 with MNE). Furthermore, CSET approximated the distribution of Fourier coefficients from the fMRI graph better than the other methods (Fig. 2e), indicating that state-of-the-art methods significantly underestimate the sparsity of brain activity in this space (KS-distance = 0.42 versus [0.59, 0.59, 0.62 0.62, 0.61]).

Qualitative assessment of the reconstructions also revealed that while the state-of-the-art methods tended to concentrate all the signal energy in frontal and parietal regions (i.e., regions close to the electrodes), CSET was able to capture ventral activation (Fig. 2f), which are notoriously difficult to capture given their distance to the electrodes.

### 3.2 Improved EEG source reconstruction accuracy

Visual evoked potentials of well-known neurophysiological paradigms (such as face or motion perception) provide data with high signal-to-noise ratio (in comparison to other types of EEG experimental paradigms) and their activation response is well documented in other imaging modalities or animal models. We reconstructed brain activity maps from the *VEPCON* EEG dataset [54] (Fig. 3a). This dataset contains two sets of recordings of visual evoked potentials: the face-stimuli dataset (response to faces vs. response to scrambled faces), and the motionstimuli (response to coherent motion vs. response to incoherent motion) (see Methods). We focus our analysis of the spatial distribution of the reconstruction on each participant’s main activation peak, i.e., the time point at which the difference in the measured EEG response between the experimental and contrast stimuli is maximal in absolute terms. As a ground truth, we compared the EEG reconstructions with the previously used fMRI response.

Figure 3 shows two improvements in reconstructing brain response during face and motion perception by the proposed CSET method over state-of-the-art methods: first, their reconstructions more accurately capture the expected activation pattern (retrieved from fMRI), as shown by higher r-squared values (Fig. 3b). Second, the reconstruction is more precise for the CSET method, as shown by the boosted signal-to-noise ratio (SNR, Fig. 3c), and the enhanced dynamics of each task’s region of interest (Fig. 3d, right fusiform gyrus (FFA) in the face task and bilateral posterior middle-temporal area (MT) in the motion task). The reconstructions are shown in Fig. 3e. The contrast between the activation levels for coherent and random motion is generally smaller than that for faces versus scrambled faces, resulting in less significant findings and a higher apparent noise level in the data for the motion stimuli.

Recognizing the potential for variability, it’s noteworthy that the optimal *λ* value might differ across individual subjects. To assess the sensitivity of our results to changes in *λ*, we conducted an analysis wherein we perturbed the optimal *λ* value by a margin of 5%. The outcomes of this sensitivity analysis, specifically focusing on the performance metrics under near-optimal *λ* ranges, are detailed in Table 1.

**TABLE 1.** Performance drop for near-optimal *λ* ranges metrics across subjects. This table summarises the distributions of percentage of performance drop (with respect to optimal performance) when varying the *λ* parameter 10% around the optimal value *λ*^∗^ ([0.95×*λ*^∗^, 1.05×*λ*^∗^]). All methods show a decline in performance with a median central tendency falling below 1%.

|  | CSET | MNE | dSPM | sLORETA | eLORETA |
| --- | --- | --- | --- | --- | --- |
| 5th Percentile | 0.19 | 0.01 | 0.01 | 0.02 | 0.01 |
| **Median** | **0.84** | **0.04** | **0.05** | **0.05** | **0.04** |
| 95th Percentile | 3.28 | 0.26 | 0.53 | 0.68 | 0.57 |

## 4 DISCUSSION

When planing an experiment requiring non-invasive neuroimaging, choosing between high temporal and high spatial resolution is a classic dilemma as usually these characteristics are mutually exclusive. Electro-magnetic tomography (ET) is in theory a promising answer to this dilemma, as it combines M/EEG recordings with structural MRI to maximize resolution in both domains. However, existing ET methods still lack accurate regularizers, resulting in limited spatial resolution and unreliable outcomes [20].

To address this issue, we propose in this article a novel ET method, Connectome Spectrum Electromagnetic Tomography (CSET), which promotes sparsity in the connectome spectrum of brain activity.

A sparse connectome spectrum implies that, at a given time point, only a limited number of eigenmodes are actively contributing to brain activity while others may have negligible contributions. The connectome Laplacian eigenmodes represent specific spatial distributions of brain connectivity that can be associated with different functional processes or cognitive functions [32]. By utilizing the eigenvectors of the Laplacian, the CSET approach leverages the brain’s intrinsic organization to identify the most relevant spatial modes that contribute to brain activity, thus helping in efficiently characterizing and understanding brain dynamics.

Based on well-established signal processing tools for solving inverse problems with signal sparsity, we incorporate diffusion MRI-derived structural connectivity data into the solution, exploiting the close relationship between brain function and brain structural connectivity [34, 35].

Previous research has explored the use of structural connectivity priors to solve the M/EEG inverse problem. While these studies have shown positive effects on reconstruction performance by enforcing smoothness among connected sources [66, 67], sparsity [68], or temporal continuity among connected sources [69], we demonstrate that our method significantly increases the accuracy and precision of reconstructed brain activity signals. Using simulated EEG signals from fMRI responses, we show that Bayesian optimization methods with brain connectivity derived regularizers capture realistic neurophysiological patterns with better accuracy than uni-modal state-of-the-art methods based on the temporal statistics of the data.

We also show that our method can reconstruct brain responses with higher spatial localization and more robustness to intrinsic noise in the EEG signal during two different perceptual processes using measured EEG signals. The signal-to-noise ratio of the reconstructed signal and the signal energy in the brain regions most involved in the task are increased. Our method takes advantage of the latest advances in graph signal processing, compressive sensing and connectomics, addressing the problems of reliability and spatial resolution simultaneously.

It is important to note that while high-density EEG systems are generally expected to offer more accurate source localization, the relationship between the number of channels and source localization accuracy isn’t necessarily straightforward (see for example [70]. Various factors, including the signal-to-noise ratio and the quality of the forward model, can influence the outcome, potentially introducing variability.

While the type of regularization is an important aspect in reconstructing electromagnetic activity, other parts of the pipeline, such as the data preprocessing, the optimization scheme, and the forward model can be equally important in influencing the reconstruction. In this work, we have not studied these other parts of the pipeline in order to focus on the inverse problem. Future studies should assess whether the performance of the proposed inverse problems is systematically affected by any steps of the pre-processing and forward modelling. In this work we have only tackled regularization in the spatial domain of the reconstruction. State-of-the-art approaches leverage the temporal smoothness of the data to enforce continuity in the reconstruction. In future approaches, we recommend tackling the combination of regularization in the spatial and temporal domains.

While using the full rank matrix U in our analysis has its advantages in preserving all available information, it is worth acknowledging that the adoption of a smaller number of eigenmodes could present a viable alternative, particularly in contexts where computational efficiency is a priority. By selecting a subset of the most significant eigenmodes, it might be possible to substantially reduce the computational cost of the algorithm without markedly compromising the integrity of the results. This approach could strike a balance between complexity and performance, allowing for quicker analyses or application to larger datasets. Future studies may explore the optimal selection criteria for eigenmodes, considering both computational efficiency and the fidelity of the underlying neurophysiological interpretation.

While the regularization parameter *λ* was tuned using a grid search, we observed that the results demonstrated relatively low sensitivity within the near-optimal range (see Table 1). Nevertheless, meticulous tuning remains essential for our problem. Future work should explore sophisticated methods that leverage the statistics of the data and/or incorporate physiological measurements. Recently, others have shown that cortical geometrical modes, derived from the cortical geometry without long-range connectivity derived, better explains the fMRI data than the connectome eigenmodes [71]. On a similar note, others have also demonstrated that is the local cortical geometry connections which plays a crucial role for the emergence of well-known functionally rellevant network harmonics [72]. These results suggest that connectome harmonics are robust to differences in long-range connectivity. This robustness indicates that our method is likely to be resilient to potential anatomical connection errors stemming from the intrinsic limitations associated with diffusion MRI data, such as missed or spurious connections.

The main limitations of assessing the performance of EEG source localization with fMRI measurements stem from the differences in their physiological sources. The distance between the EEG-generating neuronal population and the vascular supply can lead to misalignment of EEG and fMRI sources due to BOLD signal being haemodynamic. Additionally, various physiological processes requiring energetic support, such as neurotransmitter synthesis and glial cell metabolism, can cause haemodynamic BOLD changes without corresponding EEG activity. In some cases, unsynchronized electrophysiological activity or closed-source configurations may result in differential sensitivity or invisibility to EEG. Furthermore, transient electrophysiological activity may not induce significant detectable metabolic changes [73]. For the specific case of spatially localized neural activity patterns, we foresee that solving Electromagnetic Tomography by imposing sparsity the spectral graph wavelet domain using Spectral Graph Wavelet Transform (SGWT) [74] will be advantageous. Wavelets can be understood as band-pass filters on the graph spectral domain (see Fig. 1d), allowing to parameterize the signal of interest according to spatial localization and spectral scale.

In conclusion, CSET is a unique non-invasive functional neuroimaging method that offers at the same time high spatial and temporal resolution and accuracy of brain electrical activity. This is achieved by a principled combination of readily available MRI and EEG measurements. This method acknowledges the potential mismatch between fMRI and EEG sources but also leverages the statistical properties of the Bayesian approach to mitigate the risks of overfitting or artificial bias. The strategy may not resolve all discrepancies or uncertainties between fMRI and EEG, but it provides a reasoned, mathematically grounded methodology that builds upon the known function-structure connectivity relationship. It’s an evolving field, and continuous investigation and methodological refinement will be key to fully elucidate these complex interconnections. This method will enable neuroscientists to extend their current research by revisiting the already collected MRI/EEG data or plan new projects that were up to now out of reach.

## Acknowledgements

The authors would like to thank Laurel A. Rohde and Mikkel Schöttner for their helpful comments on this article.

## Funding

This work was supported by:

- Swiss National Science Foundation Sinergia Grant 170873 (PH)
- Swiss National Science Foundation Sinergia Grant PP00P1_183714 (GP)
- Swiss National Science Foundation Sinergia Grant PP00P1_190065 (GP)
- Swiss National Science Foundation Sinergia Grant PZ00P1_179988 (DP)

## Author contributions

Conceptualization: JRQ, PH; Methodology: JRQ, HF; Software: JRQ, HF, ST; Validation: JRQ, HF; Formal analysis: JRQ, HF, ST, JY, KG, PH; Investigation: JRQ, HF; Data Curation: JRQ, ST, YAG; Writing -Original Draft: JRQ, HF; Writing -Review and Editing: All; Visualization: JRQ, HF; Supervision: PH, GP; Project administration: PH; Funding acquisition: PH, GP, DP

## Conflict of interest

Authors JRQ and PH have filed a patent application related to the methods derived from this work.

## Notes

### Competing Interest Statement

Authors JRQ and PH have filed a patent application for smoe methods described in this manuscript.

### Summary of Updates

Adds changes from HBM revisions

